# The Freshwater Sounds Archive

**DOI:** 10.1101/2025.05.07.652412

**Authors:** Jack A. Greenhalgh, Mauricio S. Akmentins, Martin Boullhesen, Gabriel Lourenço Brejão, Jacob C. Bowman, Robert A. Briers, Katie Campbell, Ambrosine Clark, Mackenzie Coen, Camille Desjonquères, Soledad Gastón, Benjamin L. Gottesman, Ian T. Jones, José J. Lahoz-Monfort, Elliot Lindsay, Fernando Macías Rodríguez, Francisco Navarrete-Mier, Melissa Norton, Maria Carolina Las Casas e Novaes, Shun Okazaki, Jernej Polajnar, Milton Cezar Ribeiro, Louise Roberts, David Rothenberg, Saeed Shafiei Sabet, Ryan Satish, Brittnie Spriel, David Stanković, Kees te Velde, Jonathan H. Timperley, Katie Turlington, Jack R. Walker, Marisol P. Valverde, Kieran Cox, Audrey Looby

## Abstract

Freshwater ecosystems are full of underwater sounds produced by amphibians, aquatic arthropods, reptiles, plants, fishes, and methane bubbles escaping from the sediment. Although much headway has been made in recent years investigating the overall soundscapes of various freshwater ecosystems around the world, there remains a significant knowledge gap in our collective inability to accurately and reliably link recorded sounds with the species that produced them. Here, we present The Freshwater Sounds Archive, a new global initiative, which seeks to address this knowledge gap by collating species-specific freshwater sound recordings into a publicly available database. By means of metadata collection, we also present a snapshot of the species studied, the recording equipment, and recording parameters used by freshwater ecoacousticians globally. In total, 61 entries were submitted to the archive between the 4th of March 2023 and the 30th of April 2025, representing 16 countries and 6 continents. The most numerous taxonomic group was arthropods (29 entries), followed by fishes (14 entries), amphibians (10 entries), macrophytes (7 entries), and a freshwater mollusk (1 entry). The majority of the submissions were from European countries (27 entries), of which the United Kingdom was the most represented with 14 entries. The next most represented region was North America (11 entries), followed by South America (8 entries), Oceania and Asia (5 entries each), Africa (3 entries), and the Middle East and Central America with 1 entry each. The global south, polar regions, and areas with an elevation >500 m (asl) were underrepresented. The field of freshwater ecoacoustics to date has largely focused on the analysis of ‘sound types’ due to a current lack of knowledge of species-specific sounds. The Freshwater Sounds Archive presents an opportunity to move beyond the ‘sound type’ approach, and towards an approach with higher taxonomic resolution, ultimately resulting in species-specific descriptions. Furthermore, The Freshwater Sounds Archive will provide freshwater ecoacousticians with one of the main tools required to start creating annotated training datasets for machine learning models from soundscape recordings by referring to known species sounds present in the archive. In the long-term, this will result in the automatic detection and classification of species-specific freshwater sounds from soundscape recordings, such as indicator, invasive, and endangered species.

## Introduction

Previous research has shown that freshwater ecosystems are full of underwater sounds produced by amphibians, aquatic arthropods, (semi)aquatic mammals, reptiles, plants, fish, and methane bubbles escaping from the sediment (Decker et al., 2020; Desjonquères et al., 2019; Gottesman et al., 2020; Greenhalgh et al., 2020; 2021; 2023; Linke et al., 2018; Marian et al., 2021; Putland et al., 2020; Rountree et al 2019; te Velde et al., 2024). The soundscape, or all the sounds in an environment, comprises a combination of different sound sources: sounds produced by biological (biophony), sounds produced by climatic conditions (geophony), and sounds produced by humans (anthrophony) (Pijanowski et al., 2011; Rountree et al., 2020).

Although much headway has been made in recent years investigating the overall soundscapes of various freshwater ecosystems around the world, there remains a significant knowledge gap in our collective ability to accurately and reliably link recorded sounds with the species that produced them. The need for an archive of freshwater species-specific sounds has been identified by many authors (Linke et al., 2018; Gottesman et al, 2018; Greenhalgh et al., 2021). To address the current lack of a biological sound archive specifically dedicated to underwater sounds produced by freshwater species, here we present The Freshwater Sounds Archive (https://fishsounds.net/freshwater.js). The Freshwater Sounds Archive initiative will collate sound recordings from around the world and provide opportunities for researchers, particularly from countries with little or no current representation within the freshwater ecoacoustics literature, to collaborate and form networks with other researchers working in a diversity of freshwater ecosystems. These collaborations will help facilitate the adoption of freshwater ecoacoustics as an effective survey method more broadly within conservation biology and environmental management.

Central to the ability to understand sounds generated by the natural world is the documentation and cataloguing of sounds produced by animals, plants, and methane bubbles associated with decompositional processes. As such, the establishment and maintenance of biological sound archives is an integral part of deriving ecologically meaningful conclusions from large acoustic datasets. Since the creation of the Macaulay Library (https://www.macaulaylibrary.org/) by Cornell University, Ithaca, New York, in 1929— which is primarily dedicated to the cataloguing of bird sounds—many other biological sound libraries have been established encompassing a wide array of ecosystems and species, such as marine mammals in ocean environments (https://dosits.org/), bats in temperate woodlands (https://www.bats.org.uk/resources/sound-library), and frogs in tropical rainforests (https://www.fonozoo.com/index_eng.php) (Greenhalgh et al., 2024). Cataloguing species-specific sounds is essential if ecoacoustics is to be used in future biodiversity assessments alongside other conventional methodologies used to survey freshwater environments.

Conventionally, the ecological status of freshwater ecosystems is derived by monitoring aquatic species indicative of specific environmental conditions due to variations in their abundance, presence, or absence along an environmental gradient (Bal et al., 2018). In fact, multiple indices can be used, such as the Whalley, Hawkes, Paisley & Trigg (WHPT) index (Paisley et al., 2014), that scores aquatic invertebrates based on their preferences for varying habitat and water qualities to infer ecological status to meet legal environmental monitoring requirements, such as the Water Framework Directive (Directive 2000/60/EC, 2003). Monitoring freshwater species using conventional methods is labour-intensive, invasive, and expensive (Greenhalgh et al., 2020). Consequently, this limits the number of sites that can be surveyed, and there is a bias towards excluding hard-to-access sites from sampling campaigns that require regular visitation. Moreover, conventional methods only capture a single snapshot in time. As a result, freshwater ecologists are now supplementing conventional methods with new cutting-edge technologies, such as camera traps, environmental DNA, and passive acoustic monitoring, which involves recording all the sounds in an environment, to address ecological questions (Greenhalgh et al., 2021). Passive acoustic monitoring is rapidly becoming an affordable and effective method for non-invasively monitoring ecosystems at large spatial and temporal scales, as inexpensive acoustic sensors can be deployed for months at a time in many locations simultaneously (Browning et al., 2017; Hill et al., 2018). Additionally, passive acoustic monitoring facilitates behavioural tracking, providing insights into vital behaviours such as mating rituals, predator avoidance, and foraging (Gibb et al., 2019).

There is a wealth of ecological information that can be derived from passive acoustic monitoring of freshwater ecosystems because sounds are produced by species across multiple trophic levels and include a wide range of behaviours (e.g., mating and foraging) and processes (e.g., photosynthesis and decomposition) (Gibb et al., 2019; Greenhalgh 2023). As such, the development of the Freshwater Sounds Archive is a crucial stepping stone towards deriving meaningful ecological conclusions from passive acoustic monitoring data collected in freshwater environments. The aims of The Freshwater Sounds Archive are to: 1) Establish the world’s first global archive dedicated specifically to sounds produced underwater by freshwater species, 2) Quantitatively analyse sounds and metadata submitted to the archive, and 3) Make all the submitted recordings publicly available via the FishSounds platform to increase awareness and access.

## Methods

### Establishing the archive

Contributors to the archive were found by publishing an online form on social media platforms, and spreading the word within special working groups in bio/ecoacoustics communities for people interested in contributing. Once the initial interest had been assessed and enthusiasm for the development of an archive was made clear, collaborations with members in the community involved in running and maintaining biological sound archives followed. Namely, this involved establishing a collaboration with Audrey Looby and Kieran Cox of FishSounds (https://fishsounds.net/), an online platform that offers a comprehensive, global inventory of fish sound production research (Looby et al., 2023a). Collaboration with FishSounds therefore facilitated the future hosting of The Freshwater Sounds Archive.

A metadata file was created by JAG, AL, and KC to be filled in by all contributors submitting recordings to the archive, which can be found in the Supplementary Material (S1). Metadata parameters were decided based on their relevance for their information regarding: 1) Geography (region, country, latitude, longitude, elevation), 2) Equipment (hydrophone), 3) Recording parameters (gain, sampling rate) 4) Habitat, and 5) Taxonomy (order, family, genus, species).

### Geographical distribution

Recordings submitted to the archive via a Google Drive folder were then downloaded, along with the associated metadata. A world map was produced in QGIS (version 3.36.3) using the latitude and longitude coordinates associated with each recording and were projected onto a Google Satellite image world map. Additionally, elevation data (metres above sea level) for each recording submitted to the archive were calculated from latitude and longitude coordinates using FreeMapTools (https://www.freemaptools.com/elevation-finder.htm).

### Taxonomic representation

Taxonomic metadata were visualised in R Studio by creating a pie chart using the *ggplot2* package (Wickham et al., 2016).

### Species-specific spectrograms

Finally, species-specific spectrograms were produced in R Studio using the *spectro* function in the *Seewave* package (Sueur et al., 2008) with an FFT of 512.

## Results

### Geographical distribution

In total, 61 entries were submitted to the archive between the 4th of March 2023 and the 30th of April 2025, representing 16 countries and 6 continents (**Figure 1**). A full species list along with associated metadata is available in the Supplementary Materials (S2). The most numerous taxonomic group was arthropods (29 entries), followed by fishes (14 entries), amphibians (10 entries), macrophytes (7 entries), and a freshwater mollusk (1 entry). The majority of the submissions were from European countries (27 entries), of which the United Kingdom was the most represented with 14 entries. The next most represented region was North America (11 entries), followed by South America (8 entries), Oceania and Asia (5 entries each), Africa (3 entries), and the Middle East and Central America with 1 entry each.

**Figure 1.**
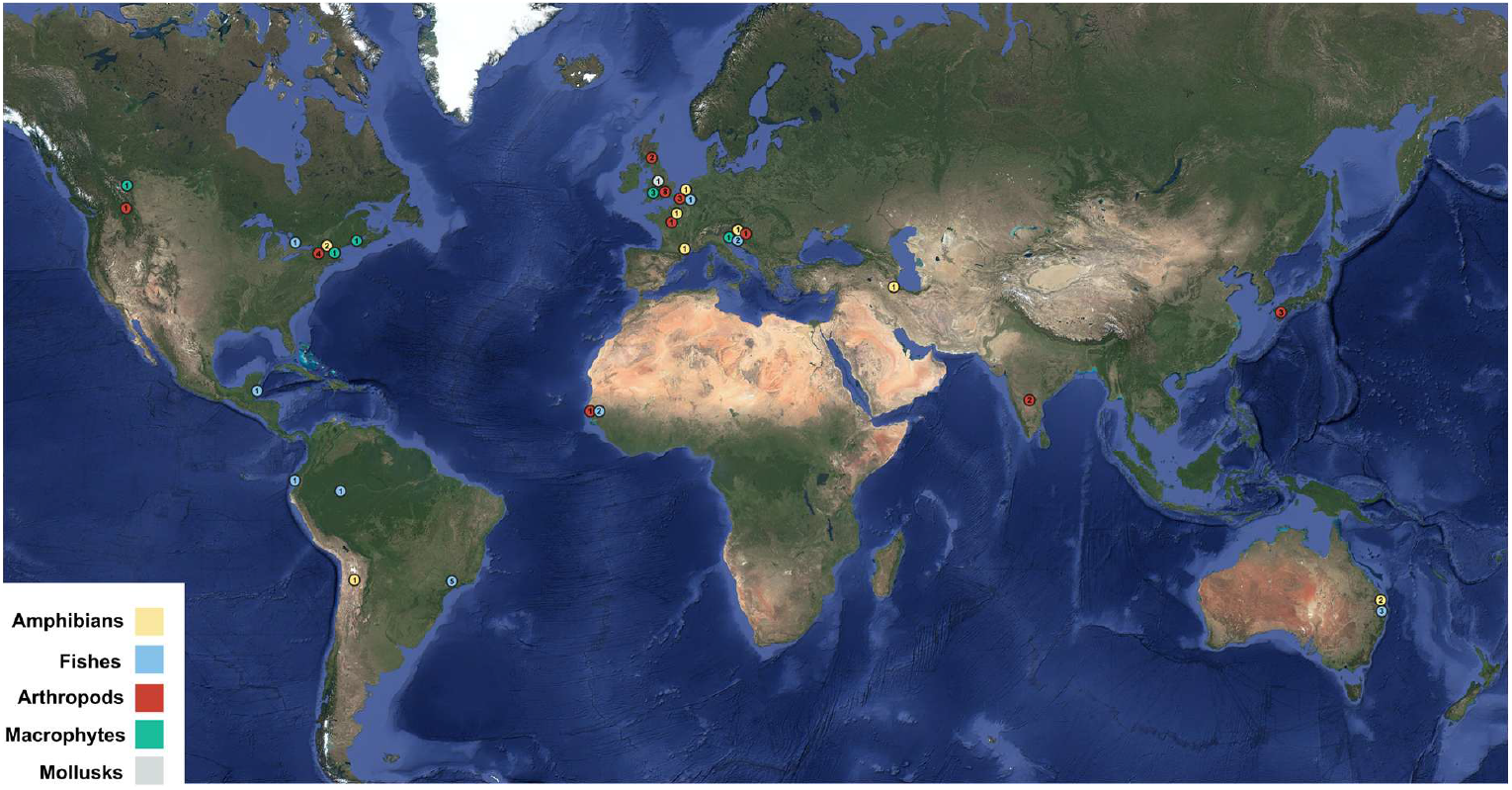
Global distribution of current submissions to The Freshwater Sounds Archive (n=61). Numbers within circles represent the number of records from each location.

The recording captured at the highest elevation was 3,797.2 m (asl) in the Argentinian Andes, and the lowest was captured at ∼0 m (asl) in an aquarium in southern Japan. The median elevation at which submitted recordings were captured was 50.3 m asl, the Q25 was 7.7 m asl, and the Q75 was 240.8 m asl. The majority of recordings submitted to the archive were collected in the northern hemisphere between 30 and 60 degrees latitude. The global south, polar regions, and areas with an elevation >500 m (asl) were underrepresented.

### Taxonomic representation

The 61 entries submitted represented 21 orders, including two unknown sounds (one likely made by a fish, and the others by plants; **Figure 2**). The most numerous taxonomic group was arthropods (29 entries), followed by fishes (14 entries), amphibians (10 entries), macrophytes (7 entries), and a freshwater mollusk (1 entry). No sounds were submitted by (semi)aquatic mammals or reptiles. In total, 4 orders and 9 different families were represented among the freshwater arthropods. The order Hemiptera (water boatmen) was the most numerous with 21 entries, followed by Coleoptera (predaceous diving beetles with 5 entries). The next most numerous group with 14 entries, fishes, was represented by 9 orders and 11 families, of which Characiformes (characins) were the most numerous with 3 entries. Amphibians were represented by one order and 7 families, of which Ranidae was the most numerous with 4 entries. Macrophytes were represented by 4 orders and 5 families. Freshwater mollusks were the least represented with one entry in one family.

**Figure 2.**
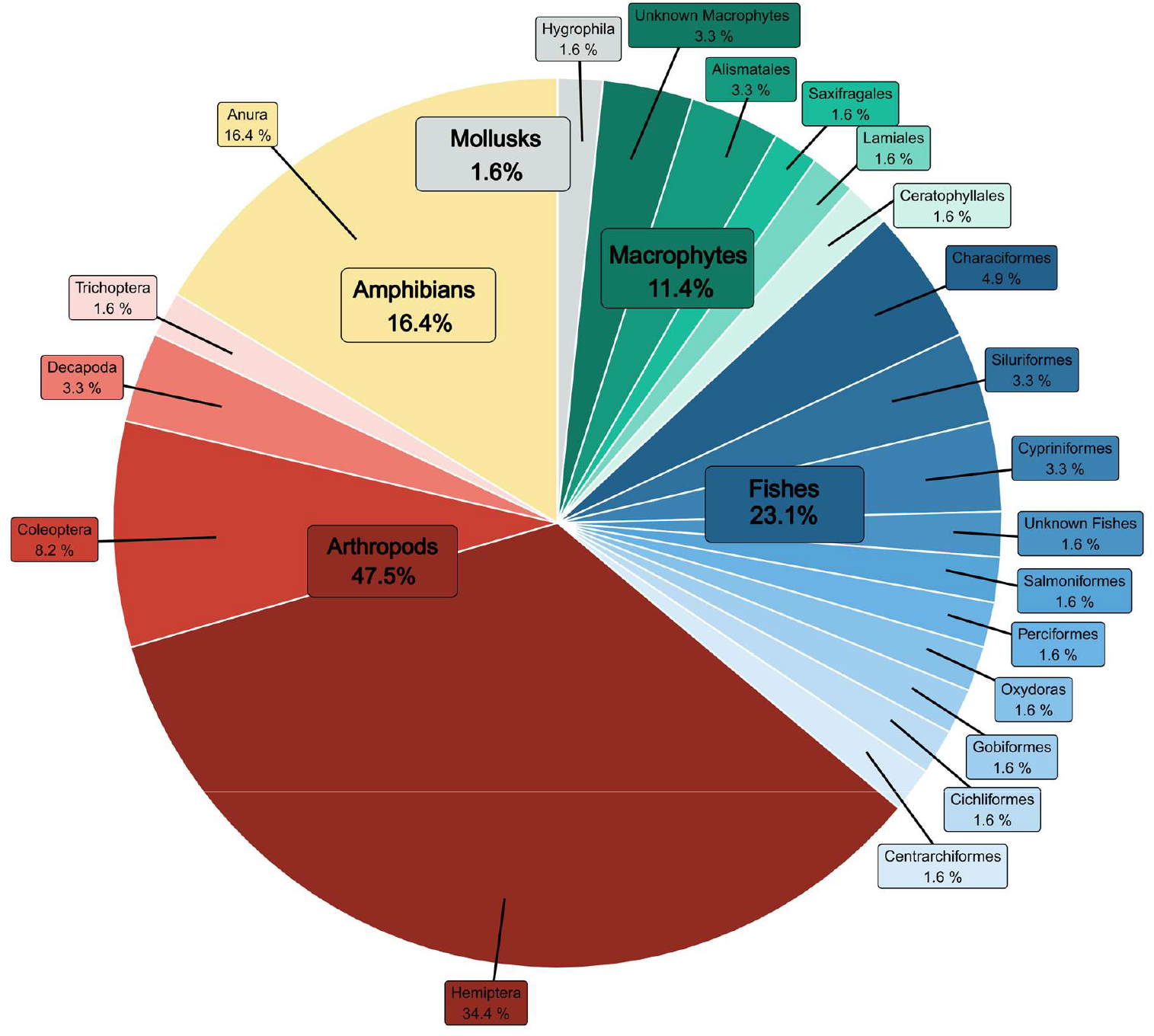
Taxonomic representation of current submissions (n=61) to The Freshwater Sounds Archive.

**Figure 2.** Taxonomic representation of current submissions (n=61) to The Freshwater Sounds Archive.

### Species-specific spectrograms

Species-specific spectrograms showed distinct patterns between taxonomic groups (**Figure 3**).

**Figure 3.**
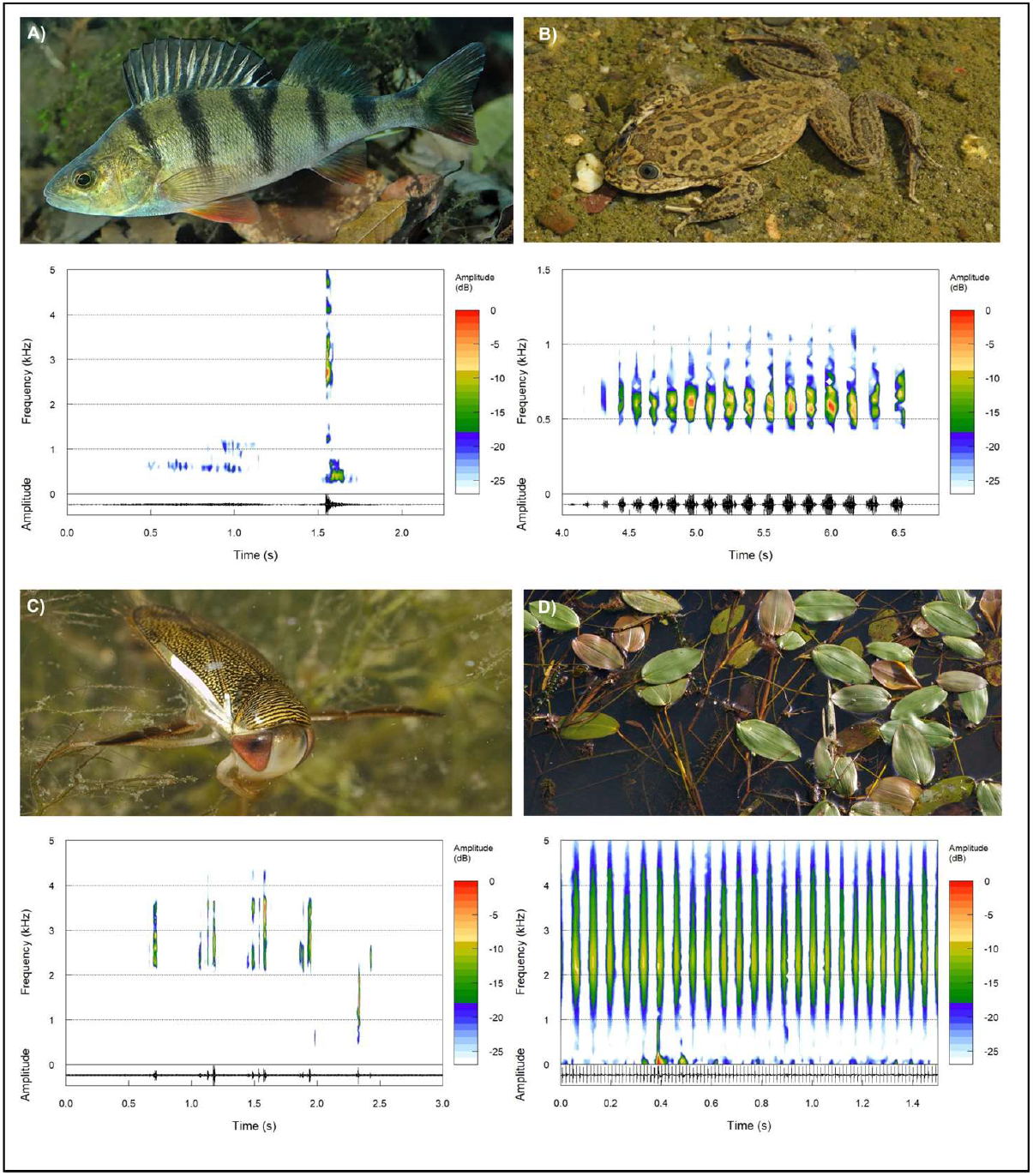
Example sounds of A) European perch (*Perca fluviatilis*), B) Rusted frog (*Telmatobius rubigo*), C) Water boatman (*Sigara concinna*), D) Broad-leaved pondweed (*Potamogeton natans*). Spectrogram parameters: window type: “Hanning”, window length: 512. Note that the frequency and time axes are not consistent across plots.

### Recording metadata

In total, 16 different hydrophones were used to collect the submitted recordings. The most popular choice was HydroMoth (12 entries), followed by the Aquarian H2a (11 entries), the HTI-96-Min (8 entries), the Aquarian H2d (6 entries), the Soundtrap 300STD (4 entries), the Aquabeat, Jez-Riley-French standard, and a custom-made hydrophone (3 entries each), the SQ26-08 Cetacean Research Tech and the Reson TC4033 (2 entries each), and the Aquarian AS-1, the Aquarian H1a, the Brüel & Kjaer Type 8103, the GoPro Hero 7, the GoPro Hero 8, and the HTI-94-SSQ (1 entry each). One entry did not disclose the hydrophone model used to collect the sounds. In total, 6 different sampling rates were used to collect the recordings. The most popular choice was 48.0 kHz (25 entries), followed by 44.1 kHz (23 entries), 96.0 kHz (10 entries), and 192.0 kHz, 45.1 kHz, and 16.0 kHz (1 entry each).

Aquaria were the most popular recording environment, with 23 entries. The next most numerous recording environment was rivers (18 entries), followed by ponds (12 entries), lakes (7 entries), and one recording that was capture in a cenote.

## Discussion

This global collaborative effort to launch the first biological sounds archive for underwater freshwater species sounds has resulted in submissions from 35 contributors represented by 16 countries and every continent except Antarctica. The establishment of a biological sound archive is essential for deriving more detailed ecological conclusions from passive acoustic monitoring data and understanding freshwater soundscape ecology.

### Describing complex and largely unknown systems

The field of freshwater ecoacoustics to date has largely focused on the analysis of ‘sound types’ (Desjonquères et al., 2015; Gottesman et al., 2020; Greenhalgh et al., 2021) due to a current lack of knowledge of species-specific sounds. In other nascent fields, similar ‘type classification’ techniques have been employed to help decode complex and largely unknown biological systems before a higher taxonomic level approach can be readily adopted. In molecular ecology, specifically environmental DNA and metabarcoding, ‘Operational Taxonomic Units’ were defined to permit the inference of taxa from unique genetic barcodes without direct reference to the species (Blaxter et al., 2005). And in remote sensing, spectral signatures (a combination of light wavelength and reflectance) are used as proxies to detect native and invasive plant species from satellite images (Iqbal et al., 2021). Whereas in complex microbial systems, in which many thousands of largely unknown bacterial and viral species are present, ecological clusters are defined as ‘phylotypes’ to begin decoding the rich biodiversity (Wu et al., 2020).

These approaches have their merits in the early development of a monitoring technique in describing largely unknown complex systems. In the field of freshwater ecoacoustics for example, the classification of ‘sound types’ has been used to make inferences of relative sound type abundance (acoustic activity) and richness across sites (Desjonquères et al., 2015), time periods (Linke et al., 2020; Gottesman et al., 2020), environmental gradients (Desjonquères et al. 2018) and management activities (Greenhalgh et al., 2021). However, this ‘sound type’ approach presents significant limitations in the ecological conclusions that can be derived due to a lack of taxonomic resolution. Furthermore, the classification of ‘sound types’ is prone to subjectivity, with sound types often receiving subjective labels such as ‘grunt’ and ‘croak’, the interpretation of which has the potential to vary between authors. It is also very likely that different sound types produced by the same species are labelled as two or more sound types, artificially increasing the derived impression of species richness (Looby et al., 2023b).

### Acoustic indices as proxies for species-specific detection

Acoustic indices—mathematical functions that consider variations in amplitude and frequency of a recording over time—have also been used to study complex soundscapes with little or no prior knowledge of the sound producers (Buxton et al., 2018). Much has been discussed in the literature on the use of acoustic indices to assess the health of coral reef soundscapes (Bertucci et al., 2016; Lamont et al., 2022), and infer avian species richness (Alcocer et al., 2022; Buxton et al., 2018; Eldridge et al., 2018). Although a suite of acoustic indices has been shown to be successful in the prediction of avian species richness where species-specific sounds are well described in the United Kingdom, they were unsuccessful in a more complex and unknown tropical rainforest soundscape in Ecuador (Eldridge et al., 2018). Moreover, a recent comprehensive review on the use of acoustic indices highlighted their limitations in inferring species diversity metrics (Alcocer et al., 2022). Soundscapes are complex systems that are influenced by many variables in addition to species diversity, such as variation in relative species abundance, vocal repertoire (number of ‘sound types’), distance from the sensor, modification by habitat components such as vegetation, and anthropogenic and natural noise, which can alter the signal to noise ratio of species-specific calls in the soundscape (Alcocer et al., 2022).

Most acoustic indices were also originally designed for the detection of bird song in terrestrial environments making them largely unsuited for the detection of aquatic fauna in freshwater ecosystems (Greenhalgh et al., 2020). Additionally, short repeating phrases that occupy a continuous frequency band with little or no variation in amplitude, such as those produced by aquatic insect stridulation, can be difficult to accurately characterise using some acoustic indices (Desjonquères et al., 2020; Ferreira et al., 2018). This is because indices like the Acoustic Complexity Index are designed to ignore consistent sounds in the recorded bandwidth to reduce the influence of anthropogenic sounds on calculated values (Pieretti et al., 2011). While some studies have successfully detected soniferous fishes and aquatic insects in freshwaters using the Acoustic Complexity Index with specially adapted parameters (Linke et al., 2020), other studies have failed to do so (Karaconstantis et al., 2020: Greenhalgh et al., 2023).

### Towards a species-specific approach

The Freshwater Sounds Archive presents an opportunity to move beyond the ‘sound type’ or acoustic indices approaches, and towards an approach with higher taxonomic resolution, ultimately resulting in species-specific descriptions. In anuran amphibians, the advertisement call is a well-recognised species-specific character used for species identification (Köhler et al., 2017). In the case of aquatic frogs, such as the genus *Telmatobius*, studies on vocal behaviour are in early stages, with some distinctions noted among species (Akmentins et al., 2024). Underwater calling represents an incipient field of work for anurans, facilitated by emergent recording technologies (Lamont et al., 2022). We have known for decades that different species of aquatic insects must be able to produce different sounds because identification guidebooks draw upon the differences in their sound-producing anatomy to distinguish between species (Savage, 1990). If different species have different anatomical structures related to sound production, then logically it follows that most must be producing species-specific sounds that can be described. Foundational work by Antti Jansson provided the first descriptions of many sounds produced by lesser water boatmen (Corixidae) (Jansson 1974, Pajunen & Huldén, 2002; Rothenberg, 2021). Multiple reviews of aquatic insect sound production since have demonstrated the widespread adoption of stridulatory behaviour by many taxa, including a recent review that estimated that more than 7,000 species of aquatic insect are likely to produce sound worldwide (Aiken 1985; Desjonquères et al., 2024). Once species-specific sounds have been identified, the large-scale annotation of soundscape recordings has the potential to generate large amounts of training data for machine learning models to automatically detect species’ sounds and potentially assess the ecological condition of freshwater habitats. The application of automated species-specific sound detection in freshwater environments has great promise in the detection of indicator, invasive, and endangered freshwater species sounds.

### Deep learning: The future of freshwater ecoacoustics analysis

One of the main challenges associated with passive acoustic monitoring is the vast amount of data that are produced, making it impossible to manually analyse recordings (Stowell, 2022). Therefore, novel computational solutions are required in the large-scale analysis of ecoacoustics data to automate large parts of the analysis workflow. Deep learning is a subset of machine learning that uses very large datasets, such as audio data collected in passive acoustic monitoring surveys, to learn from data and to make predictions using neural networks (Stowell, 2022; Dufourq et al., 2022). The most common use of deep learning techniques in ecoacoustic studies are classification and detection (Ruff et al., 2021). This is achieved by training models using pre-labelled data, such as species-specific sounds that have been annotated from soundscape recordings (Stowell, 2022). In addition, novel techniques such as transfer learning of pre-trained convolutional neural networks are promising for classification tasks (Kath et al., 2024). Biological sound archives play a crucial role in this effort by providing a starting point from which an annotated species-specific call library can be created and used to train deep learning models (Cañas et al., 2023).

Since the wider adoption of deep learning techniques within ecoacoustic studies in 2016 (Goëau et al., 2016), many authors have applied them to automatically detect species-specific sounds produced by a diversity of taxa. A recent review of the use of deep learning in ecoacoustics (Stowell, 2022) showed that of the taxa whose species-specific sounds have been studied using deep learning, birds were the most studied group (with 65 studies), followed by marine mammals (30 studies), amphibians (8 studies), bats (7 studies), arthropods (7 studies), and fishes (3 studies). However, deep learning methods are rarely used in the analysis of freshwater soundscapes. Nevertheless, Parcerisas et al., (2024) recently demonstrated that a deep learning model trained to detect underwater sound events in marine environments can transfer effectively to freshwater habitats. They further applied unsupervised clustering to identify novel ‘sound types’ based on acoustic feature similarity, a method that can aid in discovering novel sounds and looking for temporal and spatial patterns in sound events of potentially biological origin. This could be especially useful in underexplored freshwater environments. Although these results confirm that deep learning-based sound event detection works in freshwater systems, and provides a useful explorative tool, more meaningful ecological insights into species distributions and behaviours will require models trained on annotated, species-specific datasets.

As such, The Freshwater Sounds Archive will provide freshwater ecoacousticians with one of the main tools required to start creating annotated training datasets for deep learning models from soundscape recordings by referring to known species sounds present in the archive. Soon, long alphanumeric codes called Application Programming Interface keys (API keys) will facilitate the automatic downloading of annotated species-specific calls from biological sound archives for training in machine learning models to detect species-specific calls in the form of installable packages via platforms like R and Python (Stowell, 2022). API keys also facilitate data sharing between biological sound archives, which means specialised archives, such as The Freshwater Sounds Archive that mobilise a specific subset of the scientific community, can contribute to larger archives in the long-term (Scott et al., 2019). There is huge untapped potential to harness recent advances in deep learning to provide automated, scalable, and non-invasive freshwater ecological assessment. However, much work is still required to better understand species-specific sounds and build reliable annotated training data libraries.

### Challenges and limitations

In the initial stages of the archive development, it was not always possible to achieve species-level descriptions due to the challenges associated with isolating species in tanks and recording them separately. While captive auditioning has helped identify several species-specific sounds, many behaviours (e.g., mating, territory defence, distress calls) are context-dependent and less likely to occur in captivity. As such, where a species description was not possible, such as sounds submitted from soundscape recordings, an educated guess was made as to the next highest taxonomic level (at least to order level) based on physical observations of the recording environment and previous experience in listening to freshwater species’ sounds. To overcome any inaccuracies associated with this approach, all the sounds submitted to the archive will be published online via FishSounds.com and a forum will be established to permit future taxonomic revisions as more data become available and assist in the identification of new sounds. It is critically important at this early stage to avoid the mislabelling of species-specific sounds that could result in perpetuating errors throughout the wider literature. Therefore, we believe that a more conservative approach to taxonomic labelling is appropriate, unless the species has been recorded in isolation and identified by a trained professional.

## Conclusions

Passive acoustic monitoring in freshwater environments has revealed a rich diversity of sounds produced by taxa and processes that represent multiple trophic levels. This includes sounds produced by primary producers (macrophytes), primary consumers (aquatic insects), secondary consumers (fishes and amphibians), and even decompositional processes in the form of methane bubbles. Although there is great promise in the wealth of ecological data that freshwater soundscape monitoring provides, there is a considerable knowledge gap in the current inability to relate species-specific sounds with the species that produces them. The Freshwater Sound Archive therefore presents an opportunity to move past the current ‘sound type’ or acoustic index-based approach of classifying and quantifying freshwater soundscapes towards a taxonomic approach, which will lead to the inference of more meaningful ecological conclusions. Furthermore, the archive will provide freshwater ecoacousticians with the tools required to start creating annotated training datasets for machine learning models from soundscape recordings by referring to known species sounds present in the archive. In the long-term, this will result in the automatic detection and classification of species-specific freshwater sounds, such as indicator, invasive, and endangered species from soundscape recordings, as successfully achieved in other fields, such as with bird and bat species identification in woodlands, fish and marine mammals in the oceans, and frogs in rainforests.

## Supporting information

A full species list along with associated metadata is available in the Supplementary Materials

## Author contributions

**JAG:** Conceptualization; Data Curation; Investigation; Methodology; Resources; Formal Analysis; Writing: Original Draft; Writing: Review & Editing; Visualization; Supervision; Project Administration. **AL, KC**: Conceptualization; Methodology; Resources; Writing: Review & Editing; Supervision; Project Administration. **JCB, SSS, MN, FN-M, RB, AC, JT, LR, KtV, KC, MC, ITJ, JP, DS, RS, MPV, CD**: Data Curation; Investigation; Resources, Writing: Review & Editing. **DR, EL, MB, MA, SG, SO, JRW, KT, BG**: Data Curation; Investigation; Resources. **JJL-M**: Resources.

## Acknowledgements

**AL** is supported by a Banting postdoctoral fellowship. Data collection by **JCB** was supported by donations from members of the Stoney Lake community, Ontario, Canada. **MC** was supported by the National Science Foundation Research Experience for Undergraduates program: Sensors in Earth, Oceans, and Space Science, at the University of New Hampshire, U.S. **RB** was partially funded by the UK Acoustics Network (UKAN+) via EPSRC grant number EP/V007866/1. **RS** carried out data collection with support of an internal research grant at IIHS and support by Viral Joshi and Isha Bopardikar. **LR, AC, JW** and **JT** were funded by the University of Liverpool. Data collection by **MPV** was supported by a Kieckhefer Adirondack Fellowship.

## Notes

https://docs.google.com/spreadsheets/d/1O3CF10I1W2f7civewDsdPT8hqXlVdslsL3q15TMRBrI/edit?usp=sharing

### Competing Interest Statement

The authors have declared no competing interest.

